# PharmaNet: Pharmaceutical discovery with deep recurrent neural networks

**DOI:** 10.1101/2020.10.21.348441

**Authors:** Paola Ruiz Puentes, Natalia Valderrama, Cristina González, Laura Daza, Carolina Muñoz-Camargo, Juan C. Cruz, Pablo Arbeláez

## Abstract

The discovery and development of novel pharmaceuticals is an area of active research mainly due to the large investments required and long payback times. As of 2016, the development of a novel drug candidate required up to $ USD 2.6 billion in investment for only 10% rate of approval by the FDA. To help decreasing the costs associated with the process, a number of *in silico* approaches have been developed with relatively low success due to limited predicting performance. Here, we introduced a machine learning-based algorithm as an alternative for a more accurate search of new pharmacological candidates, which takes advantage of Recurrent Neural Networks (RNN) for active molecule prediction within large databases. Our approach, termed PharmaNet was implemented here to search for ligands against specific cell receptors within 102 targets of the DUD-E database, which contains 22886 active molecules. PharmaNet comprises three main phases. First, a SMILES representation of the molecule is converted into a *raw molecular image*. Second, a convolutional encoder processes the data to obtain a *fingerprint molecular image* that is finally analyzed by a Recurrent Neural Network (RNN). This approach enables precise predictions of the molecules’ target on the basis of the feature extraction, the sequence analysis and the relevant information filtered out throughout the process. Molecule Target prediction is a highly unbalanced detection problem and therefore, we propose that an adequate evaluation metric of performance is the area under the Normalized Average Precision (NAP) curve. PharmaNet largely surpasses the previous state-of-the-art method with 97.7% in the Receiver Operating Characteristic curve (ROC-AUC) and 65.5% in the NAP curve. We obtained a perfect performance for human farnesyl pyrophosphate synthase (FPPS), which is a potential target for antimicrobial and anticancer treatments. We decided to test PharmaNet for activity prediction against FPPS by searching in the CHEMBL data set. We obtained three (3) potential inhibitors that were further validated through both molecular docking and *in silico* toxicity prediction. Most importantly, one of this candidates, CHEMBL2007613, was predicted as a potential antiviral due to its involvement on the *PCDH17* pathway, which has been reported to be related to viral infections.

## Introduction

The development and subsequent market penetration of new pharmaceuticals is a critical yet time consuming and expensive process that has increased in cost by nearly 150% over the last decade. In 2016, the development of just one medicine was estimated at around $ USD 2.6 billion [1]. This is mainly attributed to the costs of pre-clinical and clinical trials where ethical issues and complications are encountered very often. As a result, only 10% of the pharmaceuticals that reach trials finally obtain FDA approval [1, 2]. For these reasons, such large investments have often limited the development of drugs for medical conditions where the niche market is not sufficient for a payback in a reasonable time frame. Even for some molecules of urgent need such as the antibiotics, where resistance is increasingly worrisome worldwide, there has been an stagnation in the discovery of alternative candidate molecules for over a decade. As a result, these issues in the discovery and production of pharmaceuticals have been seen as an opportunity to explore new approaches that combine both experimental and computational routes to accelerate the development. In this regard, some of the most successful experimental approaches include soil-dwelling, Rule of 5 (Ro5), genomics, proteomics, phenotypic screening, binding assays to identify relevant target interaction, turbidimetric solubility measurements and high throughput solubility measurements [3–7]. Despite the progress, such approaches still rely on large investments in sophisticated infrastructure for automated manipulation of samples and data collection and processing [8,9]. Alternatively, *in silico* approaches are more cost-effective and consequently, have attracted significant attention over the past few years [8,9]. Examples include virtual library screening, signature matching, molecular docking, genetic association, pathway mapping, among others [6,8,9]. In this case, however, the developed algorithms still lag behind in precision and effectiveness and the obtained candidates might require considerable experimental testing [9, 10]. This combined approach is therefore leading to the repurposing of known molecules for new and more potent treatments, which is attractive for both companies and the patients [11]. To reduce the time for screening and implementation of new therapeutic candidates even further, recent advances in artificial intelligence (AI) have provided more effective search algorithms that rely on the capacity to model relationships between the variables, which can also be trained to discover patterns in significantly large data sets simultaneously [12].

Machine Learning-based algorithms have been particularly useful for improving drug discovery because they can analyze large data sets and learn the optimal representation for specific tasks rather than using hand-craft fingerprints, which are difficult to achieve otherwise [2]. Moreover, computational techniques such as Support Vector Machines (SVM) and Random Forests (RF) have been successfully applied for the design of pharmaceuticals with high specificity and selectivity, and improved physiological behavior in terms of important parameters such as circulation times, bioavailability and biological activity [2], toxicity [13] and potential side effects [14,15]. These developments have been enabled by the availability of large public databases with information about the physicochemical and biological properties of pharmaceuticals [16]. With this information it is possible to train deep learning models, which allow virtual screening over large data sets by means of efficient optimization algorithms and new computational capabilities [17]. A recent example of the application of such models was the screening conducted by Stokes et al. [11] over a data set of more than 107 million candidate molecules. The main result was the identification of the antibiotic potential of halicin, which for the first time allowed the successful re-purpose of this molecule fully *in silico*. Halicin was originally researched for the treatment of diabetes due to the inhibition potential of the enzyme c-Jun N-terminal kinase but was abandoned because of low performance [18]. This finding provides remarkable evidence for the notion that AI is a suitable route for the screening and eventual development of new drugs. Moreover, it offers the opportunity of a reduction in both the required investment for development and the potential risks to be undertaken in pre-clinical and clinical trials. Finally, it is possible to assure that from the beginning of the development, candidate molecules comply with requirements imposed by regulatory frameworks in terms safeness and reliability.

The current global COVID-19 pandemic is a compelling example of the urgent need for automating drug discovery, as this situation is the result of a novel coronavirus (SARS-CoV-2) capable of infecting humans at an extremely fast pace [19]. To respond to this contingency, novel antiviral treatments and vaccines need to be developed in an extraordinarily short time. In this regard, according to the experts and even with the unprecedented resources allocated by governments, the shortest possible period for developing and deploying a COVID-19 vaccine is of about eighteen months [20,21]. The European Union has raised $ 8 billion for collaborative development and universal deployment of diagnostics, treatments and vaccines against SARS-CoV-2 [22]. This is also the case of the U.S. and German governments, which are planning to invest in vaccine and treatments development and distribution over $ 2 billion and $ 812 million, respectively [23,24].

Here, we applied recent developments in the field of computer vision to the critical task of active molecule prediction, which mainly involves the estimation of whether a molecule is able to bind to particular membrane receptors. Starting from the publicly available AD Dataset [25], we formalized active molecule prediction as a detection problem for which we designed an experimental framework that allowed us to evaluate results with the aid of normalized Precision-Recall curves [26]. According to our newly proposed framework, the state-of-the-art technique only performs with a 1% efficiency, however, it was reported to show an AUC score of 52% [25]. In search for a superior performance, we developed an algorithm based on deep learning for active molecule prediction, which we called PharmaNet. Our approach elevated the prediction performance (i.e., the area under the Normalized Average Precision (NAP) curve) to the unprecedented level of **65.5%**.

PharmaNet was designed on the basis of natural language processing (NLP) techniques given that in this case the most important information lies in the sequence of each of the elements. Consequently, we implemented recurrent neural networks (RNNs) as the baseline for PharmaNet due to their demonstrated performance in problems involving language [27–29]. Specifically, we considered a Gated Recurrent Unit (GRU) cell as it enables the analysis of atom sequences in an information flow direction that finalizes in the current element by analyzing the ones before it. These architectures have been used previously explored for similar tasks such as those required for property prediction and the generation of molecules according to properties of interest. For instance, Marwin et al. [30] trained Long Short-Term Memory (LSTM) cells to learn a statistical chemical language model for the generation of large sets of novel molecules with similar physicochemical properties to those in the training set. The LSTM network receives as input a canonical Simplified Molecular Input Line Entry System (SMILES) representation of the molecules. In the same way, Goh et al. [31] used SMILES as the input of a GRU to predict different chemical properties of the pharmaceuticals. In consequence, due to the versatility of the SMILES format, we implemented it for data representation in PharmaNet. This approach allowed us to build and map a 2D representation of the molecules as simple sequences of characters with varying positions in such 2D space [32].

An overview of PharmaNet information pipeline is presented in Fig. 1. We represented the input molecule as a *raw molecular image* with rows corresponding to individual atoms. Then, we processed this representation using modern visual recognition techniques. To accomplish this, we trained a convolutional encoder that gradually merges the embeddings of individual atoms with those of their neighbors, thereby resulting in a *fingerprint molecular image*. Subsequently, we input the data to a Recurrent Neural Network to analyze the information of the whole molecule and to predict a probability distribution for 102 targets. PharmaNet allowed us to classify organic compounds according to their tendency towards interaction with an active cell membrane receptor without prior knowledge of its structural features. This capability might be attractive for drug discovery approaches where the structural information of target receptors is difficult to access or non-existing.

**Fig 1.**
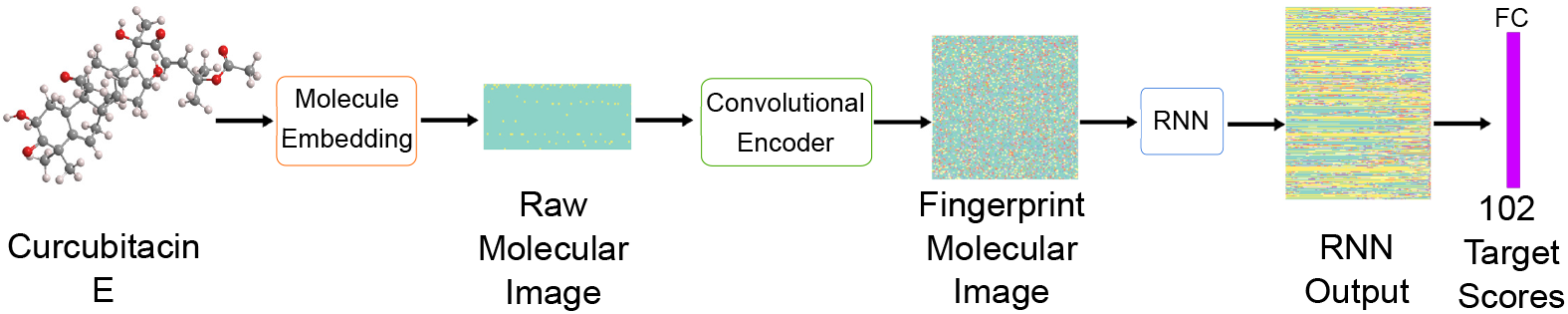
PharmaNet Workflow. For each molecule we compute a *raw molecular image* with nxm dimensions, where n is the number of unique atoms and bonds and m is the molecules’ maximum length. A convolutional encoder produces a *fingerprint molecular image* that is then analyzed globally by an RNN to predict scores for each of the targets in the AD Dataset.

PharmaNet’s architecture combines the processing power of a Convolutional Neural Network (CNN) and a RNN, which have demonstrated superior abilities to learn efficiently from image-like representations such as the proposed *raw molecular image*. Furthermore, our method enabled us to classify a molecule against multiple targets with a single trained model, which turned out to be much more efficient than most of the previously reported ones [25,33,34]. This is because as opposed to Pharmanet, such approaches are trained to classify between active and decoys for individual targets. Finally, our experiments demonstrated that all existing methods for this task provide a performance that is nominally zero, which, to our knowledge, positions PharmaNet as the most robust AI algorithm for target molecule prediction.

According to our measures, PharmaNet obtains a maximum performance for human farnesyl pyrophosphate synthase (FPPS) as a target. This protein is a key enzyme on the mevalonate (MVA) pathway that is responsible for the isoprenoid biosynthesis where it catalyzes the formation of farnesyl diphosphate (FDP). This is a precursor for several classes of essential metabolites including sterols, dolichols, carotenoids, and ubiquinones [35]. Overexpression of FPPS has been reported for multiple types of cancer, including prostate, glioblastoma, breast, and bone metastases from breast cancer. FPPS is therefore a potential target for anticancer treatments [36–41]. Also, silencing of FPPS via siRNA has slowed down viral influenza A replication and release in infected cells. This antiviral activity has been attributed to altered plasma membrane fluidity and consequently to a limited formation of the lipid rafts required for the survival of the virus [42]. On this basis, FPPS might potentially exhibit antiviral activity against enveloped viruses [42]. Given the pharmacological application for FPPS inhibitors and the high performance of our approach for FPPS, we evaluated a subset of the CHEMBL dataset to search for candidates towards this target. Our top prediction was the CHEMBL2007613 member of the data set, which corresponded to [5-[(5-amino-1H-1,2,4-triazol-3-yl)sulfamoyl]-2-chloro-4-sulfanylphenyl] acetate (PubChem CID:380934). Details of the interaction of this molecule with FPPS were determined by molecular docking analysis. Cytotoxicity and genotoxicity were further evaluated *in silico* with the aim of predicting toxicological effects directly relevant to human cells, as well as to support hazard and risk assessment activities [43]. Taken together, our results strongly suggest that CHEMBL2007613 is a potential candidate for antiviral treatments.

## Methods

### DUD-E and AD Data set

The DUD-E database contains 22886 active molecules against 102 targets and 50 decoys per target. Each decoy is a chemical compound with physicochemical properties similar to those of the active ones but different structure. Both groups of molecules (i.e., active compounds and decoys) have therefore different data distribution, thereby making the binary classification problem more amenable for a neural network. This approach has been proved successful previously by Gonczarek A. et al. [34] and Chen et al. [25].

The Active Decoys (AD) data set was proposed by [25] as a strategy to eliminate the bias introduced by the decoys. This data set is based on DUD-E but changes the decoys of each target by those contained within the 101 receptors with the highest affinity towards the target as estimated by molecular docking. This approach leads to a rather challenging binary classification problem because all the molecules show the same data distribution.

### Data Preparation

The output of our model for a molecule is the binding probability distribution to individual targets within the set of target classes. This was accomplished by defining a multiclass classification problem instead of a binary one. For this reason, the ligand sequence is labeled with the corresponding target protein prior to be input into our model.

We randomly split the complete set of binding ligand-protein SMILES sequences from the AD data set into two main subsets where 90% of the available active molecules for a target helped training the model, while the remaining 10% were employed for testing purposes. Then, we split the training subset into four folders by making sure that all the folders had the same distribution of active molecules per target. Subsequently, we conducted a four-fold cross-validation approach where data in three of the folders were used to train each model while the remaining one was only for validation. This multi-step validation approach allowed us to train the model very robustly and finally report the ensemble of the four models as an overall metric. Lastly, we converted the SMILES representation of the active molecules in the data set into *Raw molecular images*. This process is described into detail below.

### Neural Network Design: PharmaNet

PharmaNet comprises three main phases (Fig. 1). In the first one, a SMILES representation of the molecule undergoes an embedding stage that results in what we denominate a *raw molecular image*. In the second phase, that representation is the input of a CNN to obtain a *fingerprint molecular image* that incorporates information of each atom and its neighbors [44]. This stage allows the model to extract fingerprints of functional groups in the molecule, which are essential in defining their functionality. After this stage, the CNN’s output is processed by an RNN architecture that enables a sequence-based analysis of the molecules [27]. This approach enables precise predictions of the molecules’ target on the basis of the feature extraction, the sequence analysis and the relevant information filtered in the last two stages. The extracted information allows a different representation of the molecule in which the model can learn the features that determine the activity towards a target protein. For this purpose, we compute the probability scores from the RNN’s output with the aid of a Fully Connected (FC) layer, followed by a Softmax Classifier.

### Raw Molecular Image

PharmaNet’s input is a SMILE sequence embedded into a one hot vector as performed in [31]. We then generated a matrix of size n x m, where n = 36 is the total of unique chars (atoms and type of bonds) in the data set, and m = 116 is the longest SMILES sequence over all the molecules. This representation can be seen as a two dimensional image of the molecule. This type of data arrays are typically encountered in computer vision algorithms, which is attractive as we have extensive expertise with such approaches [45–47]. As delineated below, we indeed applied our most recent developments and insights in classification algorithms to classify them accurately.

### Fingerprint Molecular Image

As shown in Fig. 1, the *raw molecular image* enters a feature extracting phase. The image passed through a convolutional encoder composed by two 2D convolutional layers, ReLU nonlinearity activations and batch normalizations. By following ResNet’s central concept [48], we performed a residual connection between the convolutional layers. For the first layer, we computed the feature maps with a 5×5 kernel to produce a 64 channels output. The second convolutional layer used a 5×5 kernel to produce 128 output channels. The final output is a matrix representation of each atom that takes into account several of its neighbors. The obtained matrix can be interpreted as a *fingerprint molecular image* of functional groups for the chemical compound.

### Global analysis by recurrence

The *fingerprint molecular image* was input to an RNN, which is a bidirectional Gatted Recurrent Unit (GRU) of 10 layers. This RNN analyzed the sequence of atoms by selecting the information flow from the current fingerprint with that of the atoms previously considered. Each GRU’s cell comprises two gates that control the flow of information between them, namely, the reset gate (*r*) and the update gate (*z*). (*r*) decides whether to keep, in the current cell state (*h’*_*t*_), information of the previous cell or to change it by information from the input (*x*_*t*_). The recurrence of this process is described by the following set of equations:

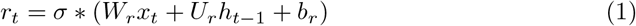

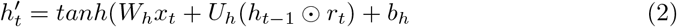

In parallel, (*z*) controls which type of information of the previous and current cell will go to the next one. The process is described by the following set of equations

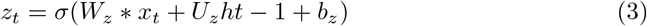

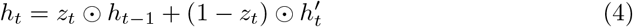

where W and U are weight matrices, b is a non-linearity, h’ is the current cell state and h is the output of the memory block [27,49]. Finally, we considered the last hidden state of the GRU, which contains information of each atom based on both its neighbors atom type and its position in the SMILES. This output is processed by a FC layer and a Softmax Classifier to compute the probability distribution over the 102 target receptors. This multiclass classification task was accomplished with the aid of a Cross Entropy Loss.

### Implementation details

The training curriculum was implemented via an Adam optimizer with a starting learning rate of 5*x*10^*−*4^, betas of 0.9 and 0.999, an epsilon of 1*x*10^*−*8^ and a weight decay of 0.1. We reduced the learning rate when the validation loss stagnated at both a factor of 0.1 and a patience of 10. We trained the model end-to-end during 30 epochs with a batch size of 128. For the network, we implemented a four-fold cross-validation approach.

### Evaluation Metrics

In order to measure algorithmic performance, both the DUD-E and AD data sets report the area under the Receiver Operating Characteristic curve (ROC-AUC). The ROC curves feature Fallout against Recall, where Fallout (F) is the probability that a true negative was labeled as a false positive while Recall (R) is the fraction of the true positives that are detected rather than missed [50]. Considering that we are classifying molecules according to their binding receptor but against 101 no binding receptors, we can understand our problem as a highly unbalanced detection task. Consequently, when evaluating a single target, negative samples largely exceed the positives ones. As a result, if we evaluate in light of the ROC-AUC curves, the probability of predicting undesirable true negatives increases significantly. In this scenario, Precision-Recall (PR) curves represent an attractive alternative to ROC curves because detection tasks proceed by normalizing Precision (P) over the true positives [51–53]. The area under the PR curves has been typically reported as the main metric for detection problems, which has been also known as Average Precision (AP).

Nevertheless, as pointed out by Hoiem et al. [26], AP strongly depends on the true positive samples in each class t (*N*_*t*_). As a result, the best performance is for classes with the largest numbers of true positives. To create a Normalized Precision *P*_*N*_, [26] replaces (*N*_*t*_) with a constant *N* that corresponds to the average of positive samples over all the classes. Eq. 5 corresponds to the definition of Normalized Precision (P_N_), where R is the fraction of objects detected while F is the number of those incorrectly detected.

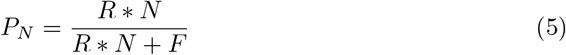

As not all targets have the same number of active molecules, we prefer to use *AP*_*n*_ for evaluating the prediction on each target. We tested the four trained models in the test subset and computed the *AP*_*n*_ for each target. With the estimates of mean and standard deviation for the four models, we then computed the Normalized Averaged Precision (NAP) curve based on the frequency of targets that achieve each *AP*_*n*_ score. We estimated the overall performance of PharmaNet as the area under the NAP curve normalized by the total number of targets.

### Screening in ChEMBL

We evaluated our best trained model with the ChEMBL database [54]. This manually curated database contains information on chemical, bioactivity and genomic data of over 15’504.604 bioactive molecules with drug-like properties [54]. Within ChEMBL, 15’367.297 members have their chemical structures represented by SMILES. This large volume of information requires a wide search field to predict the activity of ChEMBL molecules towards the 102 receptors in the AD data set.

Within ChEMBL, we only selected the molecules with lengths below those of the largest SMILE in the AD data set. Also, those exhibiting the same components and SMILES representations in AD data set. After applying this filter, the screening was reduced to 13’827.575 SMILES towards 102 targets. We computed the probability of each molecule to be a binding-ligand for each target, and predicted whether it was active towards the one with the highest score. Then, we focused on the molecules predicted as active toward FPPS and sort them out to find the top-10 compounds for which the network was more confident at the prediction.

### Molecular Docking

The top 10 candidates for human farnesyl pyrophosphate synthase (FPPS) predicted by PharmaNet were further corroborated by molecular docking via AutoDock 4.2.

### Target Protein

The target protein in this study was the human farnesyl diphosphate synthase. The 3D structure for this protein was downloaded from Protein Data Bank Database (PDB ID: 1ZW5). Protein optimization was performed by removing all water molecules and all other molecules outside of the A chain of the target protein. Polar hydrogen groups were added, and finally, Kollman charges were computed. A PBDQT file was produced using MGL-Tools in AutoDock.

### Ligand Preparation

SDF files of candidates were downloaded from PubChem and converted to PDB files with the aid of PyMOL. Ligands were optimized and converted to PDBQT files via the MGL-Tools in AutoDock. For optimization Gasteiger charges were assigned and non-polar hydrogens were combined. The rigid roots of each compound were defined automatically and rotation of amide bonds was fully restricted.

### Molecular Docking Parameters

Molecular docking on Zoladronate’s binding site was performed by following [55]. The grid box size was set at 40, 40, and 40 Å(x, y, and z) and Auto Grid 4.2 in conjunction with Auto Dock 4.2 were used to produce the grid maps for each ligand. The distance between grid points was 0.375 Å. To search the best conformer, the Lamarckian Genetic Algorithm was implemented with a maximum of 10 conformers per ligand. The size of the population was set to 150 and the individuals were initialized arbitrarily. The maximum number of energy estimation was set to 2500000 while the maximum number of generations to 27000. Also, the maximum number of top individuals that automatically survived was set to 1 with a mutation rate of 0.02, a crossover rate of 0.8. The step sizes were 0.2 Åfor translations, 5.0^*°*^ for quaternions and 5.0^*°*^ for torsions. Cluster tolerance was maintained at 2.0 Åwith an external grid energy of 1000.0 and a max initial energy of 0.0. The max number of retries was set to 10000 for 10 LGA runs. The highest binding energy (most negative ΔG) was considered as the ligand with maximum binding affinity. A positive control of the binding algorithm was the well studied inhibitor for FPPS, Zoladronate [55].

### Verification of AD data set distribution

In order to verify the complexity of the binding ligand-receptor task in the AD data set, we applied molecular docking for the best five hundred (500) molecules predicted by PharmaNet for each target. This data set encompassed both active and inactive molecules for the receptors. We used the binding energy as classification -parameter-by multiplying the energy by −1 and computing the corresponding sigmoid function. The obtained data was interpreted as the classification probability for active molecules and allowed us to compute the precision recall curve over all the classes and the mean average precision.

### Toxicity evaluation *in silico*

The top 3 candidates obtained by molecular docking were further evaluated for cytotoxicity and genotoxicity via the on-line Way2Drug Predictive Services server [56]. Citotoxicity was performed using the SDF file of the ligands as input to the GUSAR Software. This software predicts the LD50 for four routes of pharmaceutical administration (intraperitoneal, intravenous, oral and subcutaneous) and calculates a toxicity class based on the OECD principles [57]. Genotoxicity evaluation was conducted with the aid of the DIGEP-Pred software. This tool is a web-based service for *in silico* prediction of drug-induced changes of gene expression profiles based on the structural formula of the compounds [58]. This server received as input the SMILES sequence of the ligands.

## Results and Discussion

### Verification of AD data set

We performed molecular docking for 51000 molecules and found that 1926 of them showed activity towards one of the 102 targets. With the binding energy, we calculated the probability for a binary classification and computed the average precision (AP) of these predictions. Fig. 2 shows the precision recall curve for the active classes with an area under curve, i.e., AP of 3.8%. This result indicates that classification by molecular docking essentially leads to the same result obtained by random classification.

**Fig 2.**
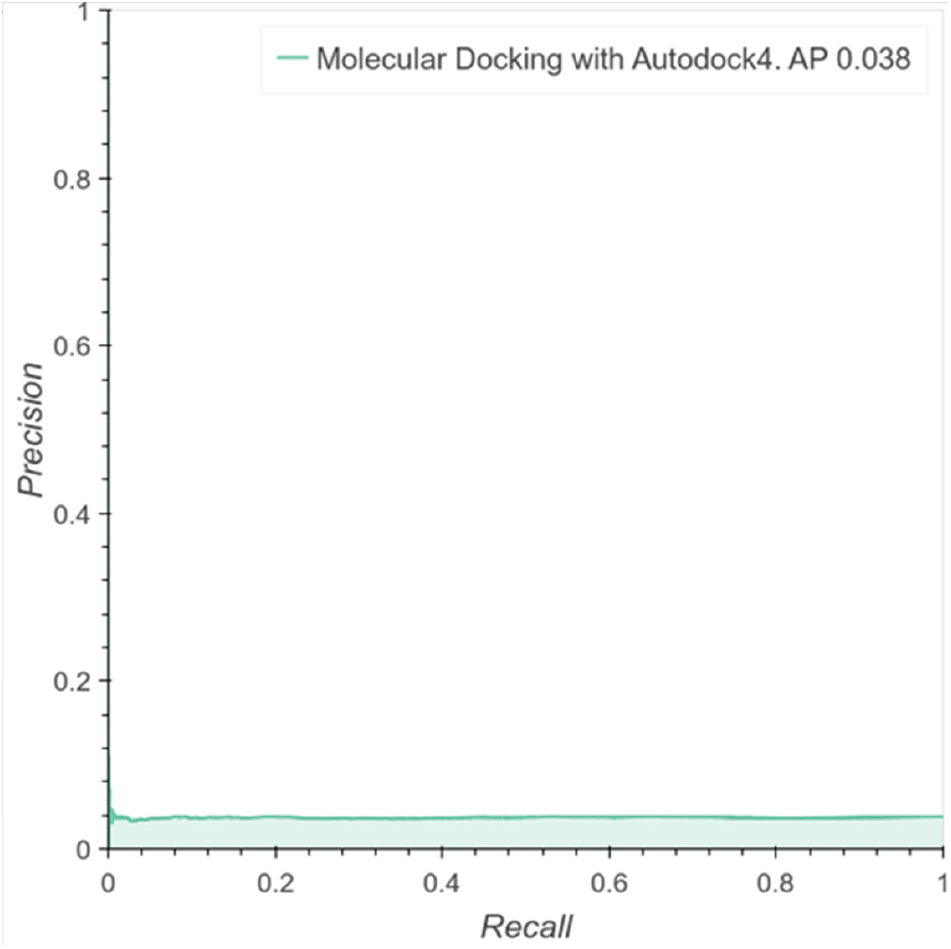
Performance of Molecular Docking as a Classifier. Precision Recall Curve over the predictions computed with the binding energy of the molecular docking in the selected molecules against one of the 102 targets.

Furthermore, this experiment confirms the need for new computational tools for more robust and accurate analyses of scenarios where active and inactive molecules toward some targets have similar data distributions, molecular structures, length and physicochemical properties.

### Comparison with the state-of-the-art AI algorithms

We compared the performance of PharmaNet against the prediction made by the state-of-the-art method recently published by Chen et al. [25] for each target in the AD data set. The data was kindly provided by the authors for the comparisons presented herein.

Fig. 3A shows the behavior of Chen et al.’s method [25] for the 102 targets classes evaluated, where half of the models separate active molecules and decoys with an AUC performance of 51%. After reaching this point, their curve decreases rapidly to zero to reach an overall performance of 51.8% (Curve Area in the AUC frequency curve).

**Fig 3.**
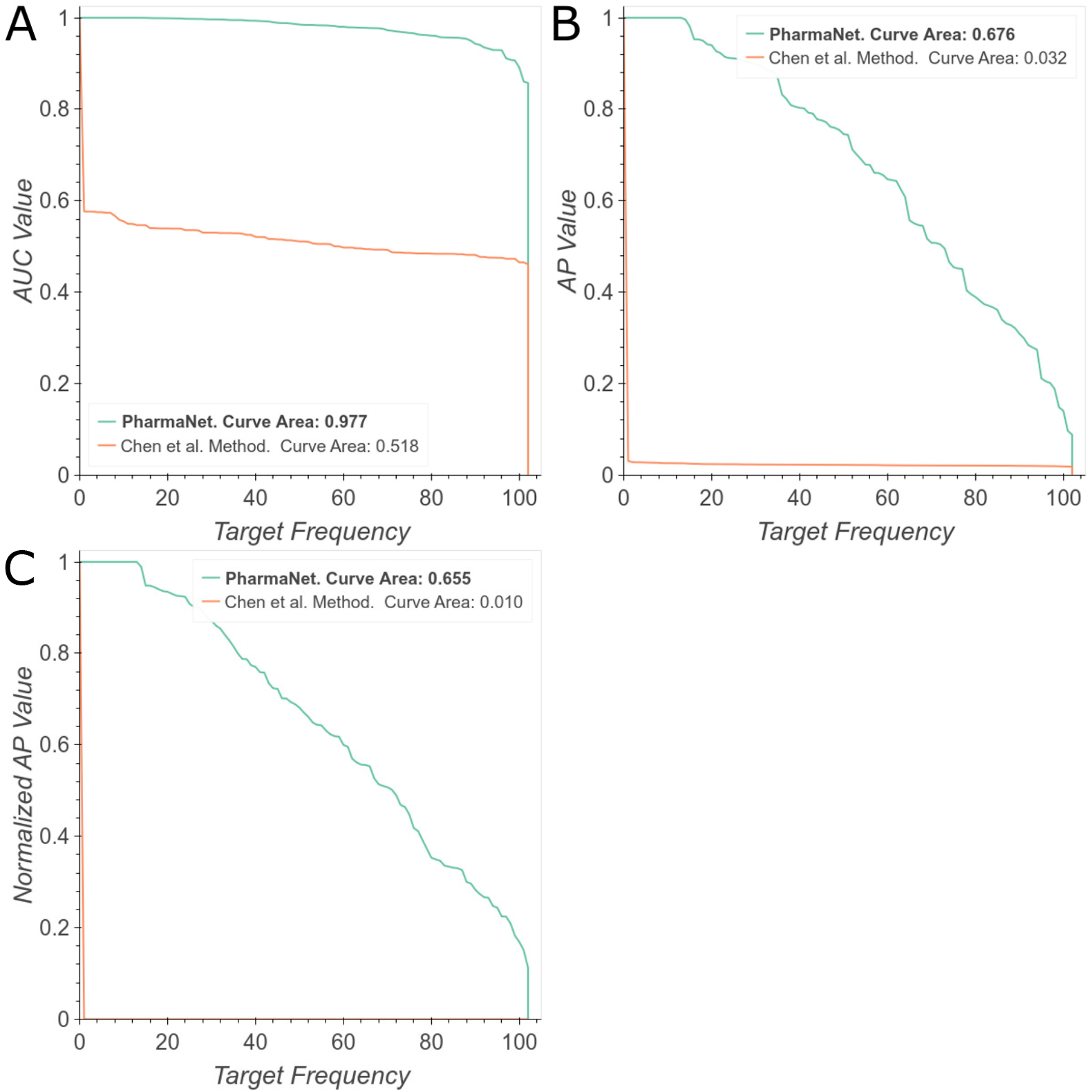
Comparison of PharmaNet against the state-of-the-art method of [25]. The curves correspond to the frequency in the 102 targets for each metric score: (A) Area Under the ROC Curve, (B) Average Precision, (C) Normalized Average Precision. We report area under the frequency curve. Best viewed in color.

While [25] trains a different model for each of the 102 target molecules, our single model is able to predict targets with a general performance of 97.7%. However, a relatively small decrease in performance was observed after 0.8. A more stringent evaluation shows an area under the AP curve of 3.2% for Chen et al.’s method [25] while for PharmaNet is 67.6% (Fig. 3B). We obtained consistent results when we evaluated both methods with the area under the NAP curve, in which [25] achieved 1% while ours approached to 65.5% (Fig. 3C).

### Ablation Experiments

To verify whether all the components of PharmaNet were relevant to the task and to select optimal hyperparameters, we performed an ablation study. First, we evaluated the main phases of the architecture and their performance in the prediction task. Fig. 4A shows the configuration of PharmaNet for our best result, the same architecture without the convolutional encoder and without the RNN. It is evident that the RNN is the most important component for superior performance but their combination is still beneficial for improving performance somewhat further. Regarding the specific RNN implemented herein, Fig. 4B shows that training with an LSTM decreases significantly PharmaNet’s performance. Considering the amount of parameters that an LSTM has to learn and the time to train this network, we decided to keep the GRU as our main RNN architecture.

**Fig 4.**
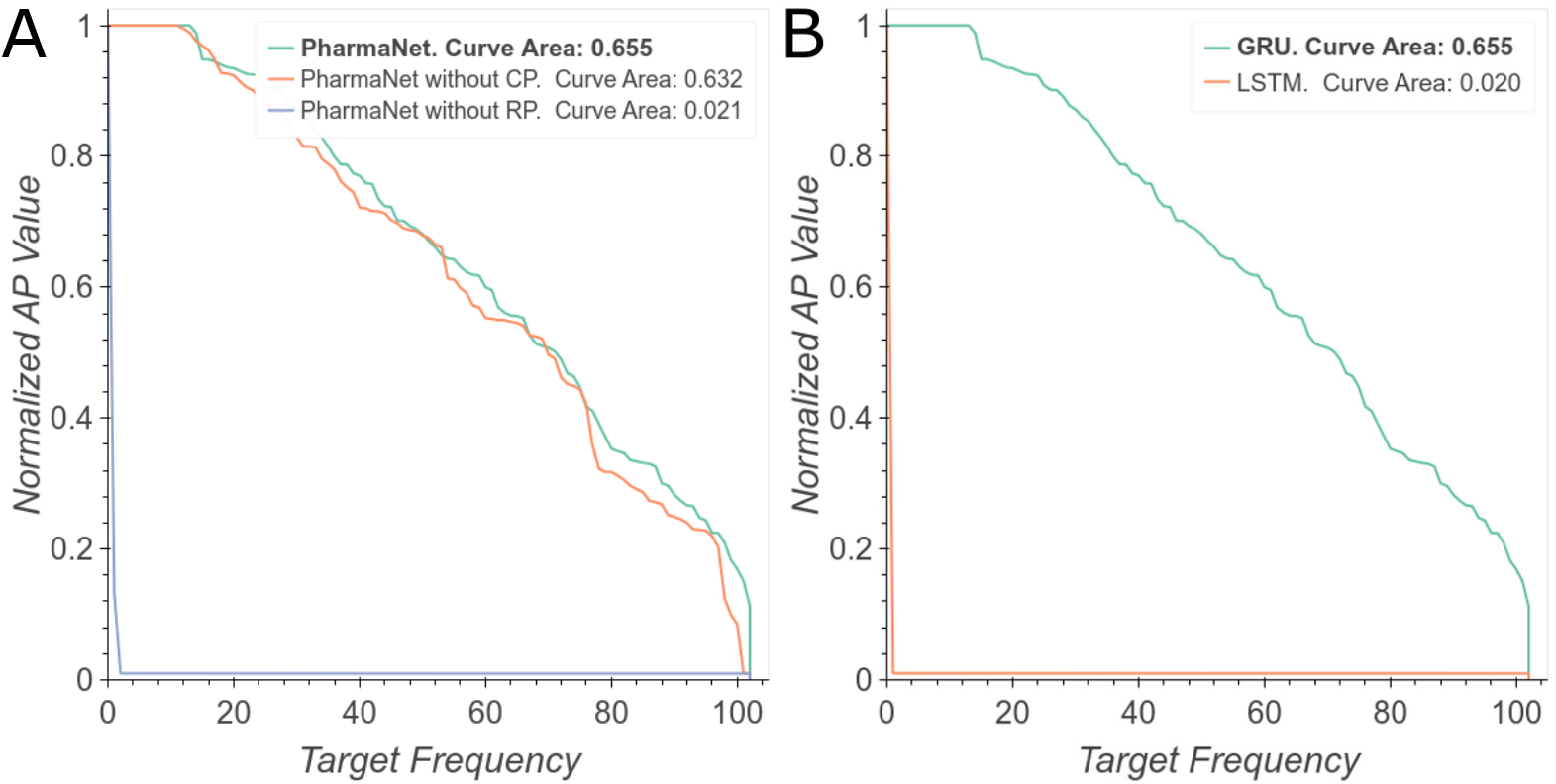
Main ablation experiments. (A) NAP curves evaluating the three main phases of our architecture. (B) NAP curves evaluating different RNNs. Best viewed in color.

Regarding the Convolutional Encoder, experiments show that the best performance was achieved with two convolutional layers and a residual connection (Fig. 5A, and 5D). However, we obtained similar results after eliminating one layer. We also found that batch normalization produced the best results (Fig. 5C) and that the optimal kernel size was 5×5 (Fig. 5B).

**Fig 5.**
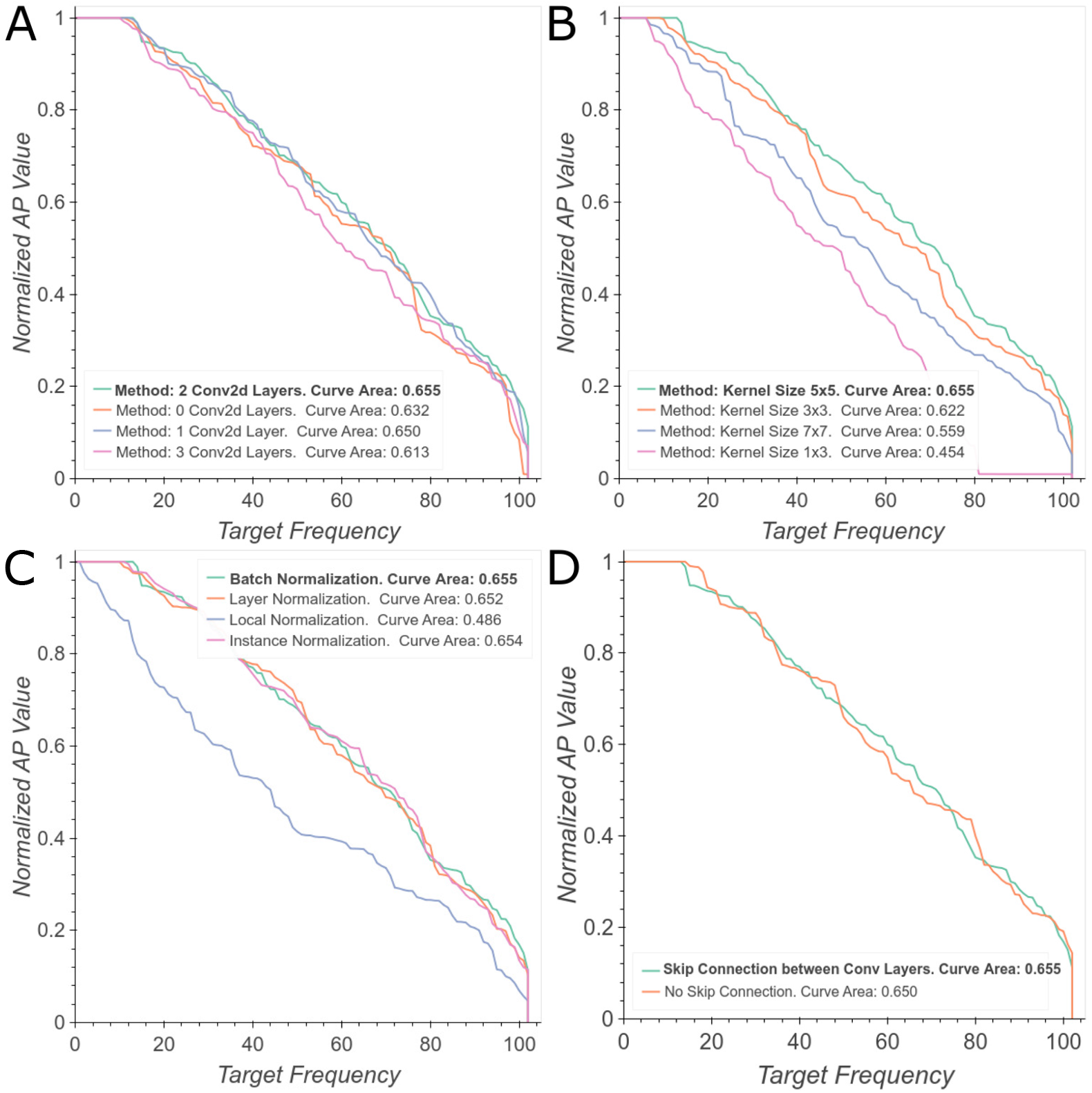
Ablation study of Convolutional architecture. (A) Number of convolutional layers. (B) Kernel size for the convolution. (C) Type of normalization layers. (D) Type of residual connection. Best viewed in color.

We also studied directionality, depth and hidden state size to establish the best configuration for the RNN. Fig. 6A shows that an bidirectional configuration leads to an slight improvement in performance compared with unidirectional GRU cells. This result is obtained by the analysis of the molecule from both left to right and right to left most likely due to the complementarity of the fingerprints obtained by the analysis of each direction. Regarding deepness of the network, Fig. 6C shows a significant increase in performance from 0 to 10 layers followed by a decrease afterwards. Finally, Fig. 6B demonstrates that by increasing the hidden size from 116 to 256, the performance increase in about 40%.

**Fig 6.**
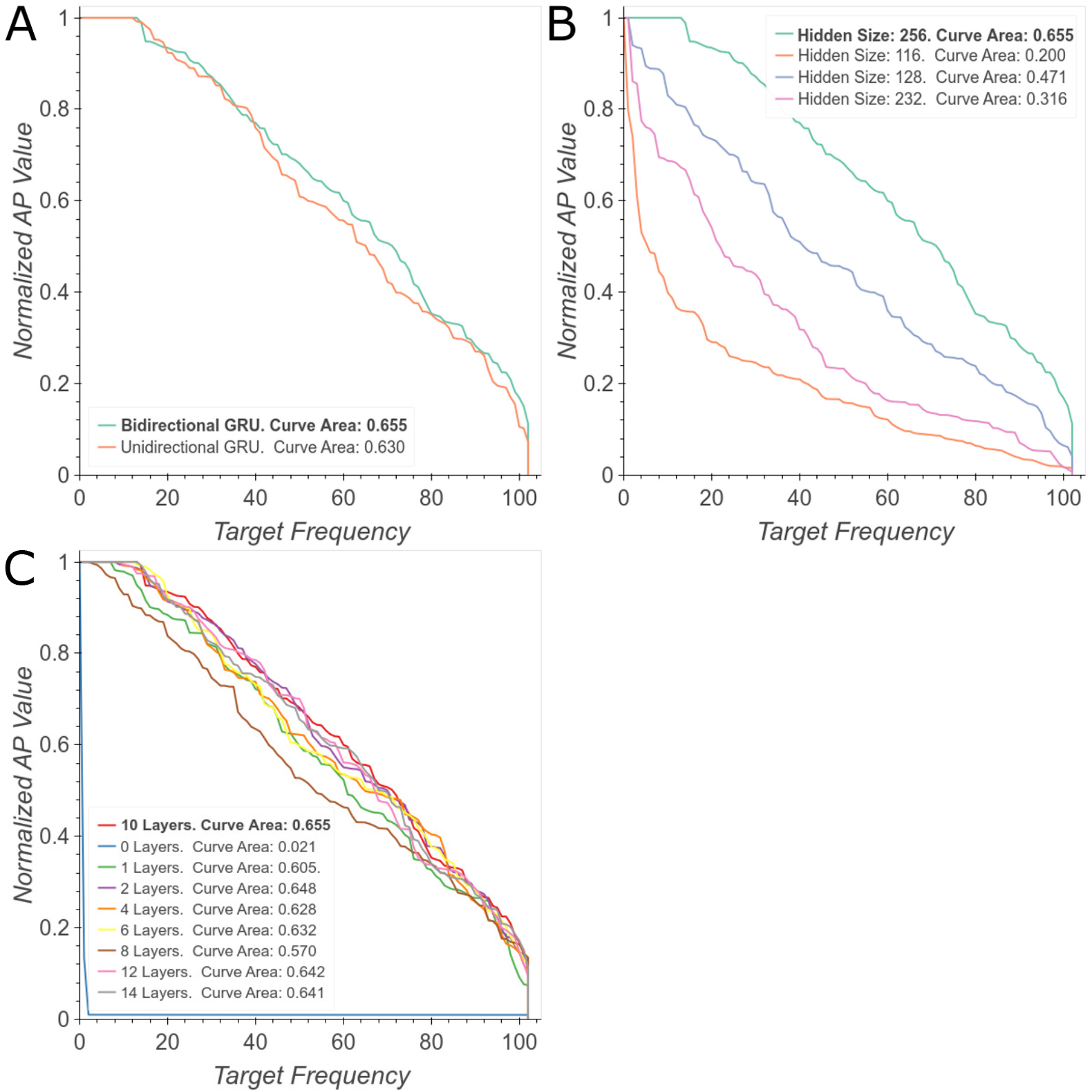
Ablation study of GRU’s architecture. (A) Unidirectional vs. Bidirectional GRU. (B) Hidden State Size. (C) GRU’s depth. Best viewed in color.

### Targets Analysis

Table 1 summarizes the performance and possible pharmaceutical applications of the top-10 best-performing receptors. Most of these receptors exhibit potential antimicrobial activities, which is attractive not only to pharmaceutical industries but also to public health, giving the increasing resistance of microorganisms to conventional antibiotics [59]. In this regard, having a method to predict with high accuracy new active molecules towards the target proteins might propel the drug discovery process unprecedentedly. Also, we identified multiple possible targets for the development of novel pharmacological anticancer therapies. This is particularly important for certain types of cancer that are resistant to conventional chemotherapy such as doxorubicin [60], imatinib [61], nilotinib [62], cisplatin [63], tamoxifen [64], paclitaxel [65], temozolomide [66], and docetaxel [67]. Finally, one of the receptors could be a potential target for Parkinson’s disease and consequently, a route to improve the palliative treatments for the disease.

**Table 1.**
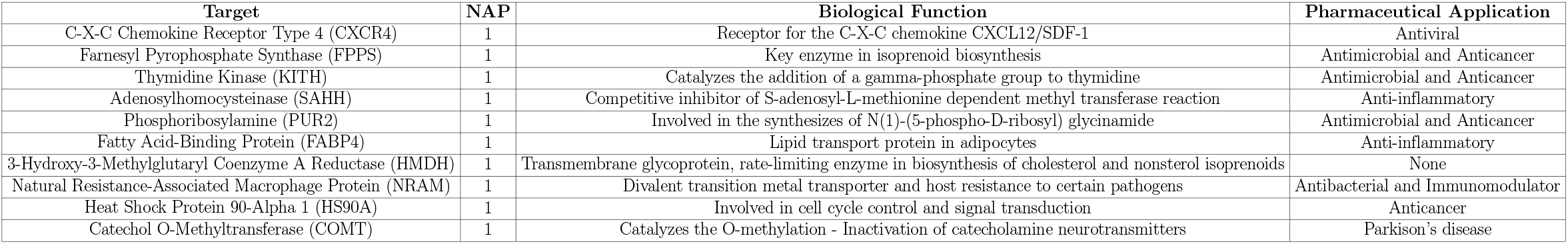
Name, biological function and pharmaceutical application of best 10 performing targets with PharmaNet.

A closer inspection of each class separately allowed us to identify certain functional groups that enabled a better classification, as we illustrate in Table 2.

**Table 2.**
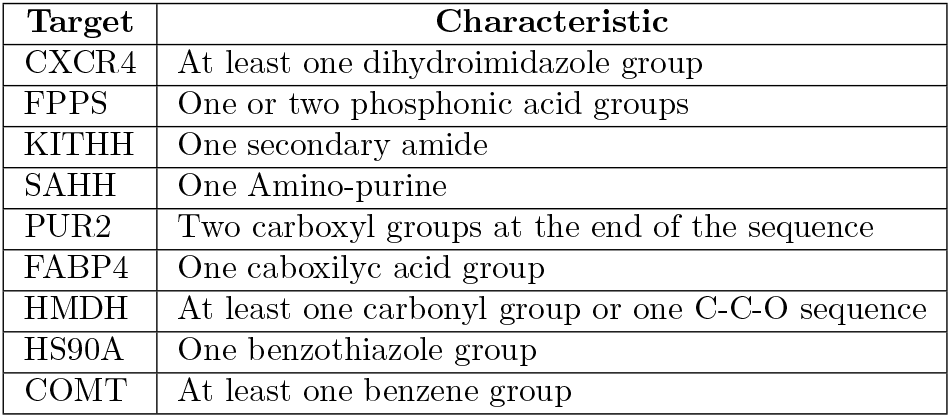
Chemical characteristics of the best performing classes with PharmaNet.

| Target | Characteristic |
| --- | --- |
| CXCR4 | At least one dihydroimidazole group |
| FPPS | One or two phosphonic acid groups |
| KITHH | One secondary amide |
| SAHH | One Amino-purine |
| PUR2 | Two carboxyl groups at the end of the sequence |
| FABP4 | One caboxilic acid group |
| HMDH | At least one carbonyl group or one C-C-O sequence |
| HS90A | One benzothiazole group |
| COMT | At least one benzene group |

The top-6 worst-performing classes were ABL1, CP3A4, CP2C9, SRC, LCK, and CDK2. Their active molecules, however, vary widely within each class, both in length and in the type of present functional groups. These organic molecules are composed principally by chains of C, N, O, H, and contain from few to none P, Cl, Br, I, S, and Si in their structures. The process of learning functional fingerprints was relatively hard for the network due to the similarities between the involved molecules. Furthermore, the activity towards the ligands for some protein subclasses might vary significantly due to 3D changes in their binding pocket as a result of the co-existence of multiple conformers. This is actually the case of the ABL1 protein subclass. Considering such heterogeneity and the corresponding volume of data, our network is unable to learn one representation that can be generalized across all the different types of ligands for making meaningful predictions regarding their activity towards different conformers of the same protein

To verify that the learning process of our network involved no simple similarity correlations among the active molecules for a specific target, we calculated the performance of our model per protein class as a function of the Tanimoto Coefficient (TC). We employed RDKit to compute the Morgan Fingerprints of the molecules to subsequently calculate their Tanimoto similarity. We averaged the TC of the molecules with the same target class and plotted these values against PharmaNet’s results per protein (Fig. 7). We also computed a coefficient of determination (*R*^2^) of 0.425, which strongly indicates a low correlation between the TC and the Normalized AP per protein. This result therefore suggests that a mere extrapolation of TC is not enough to obtain suitable candidates for the targets of interest.

**Fig 7.**
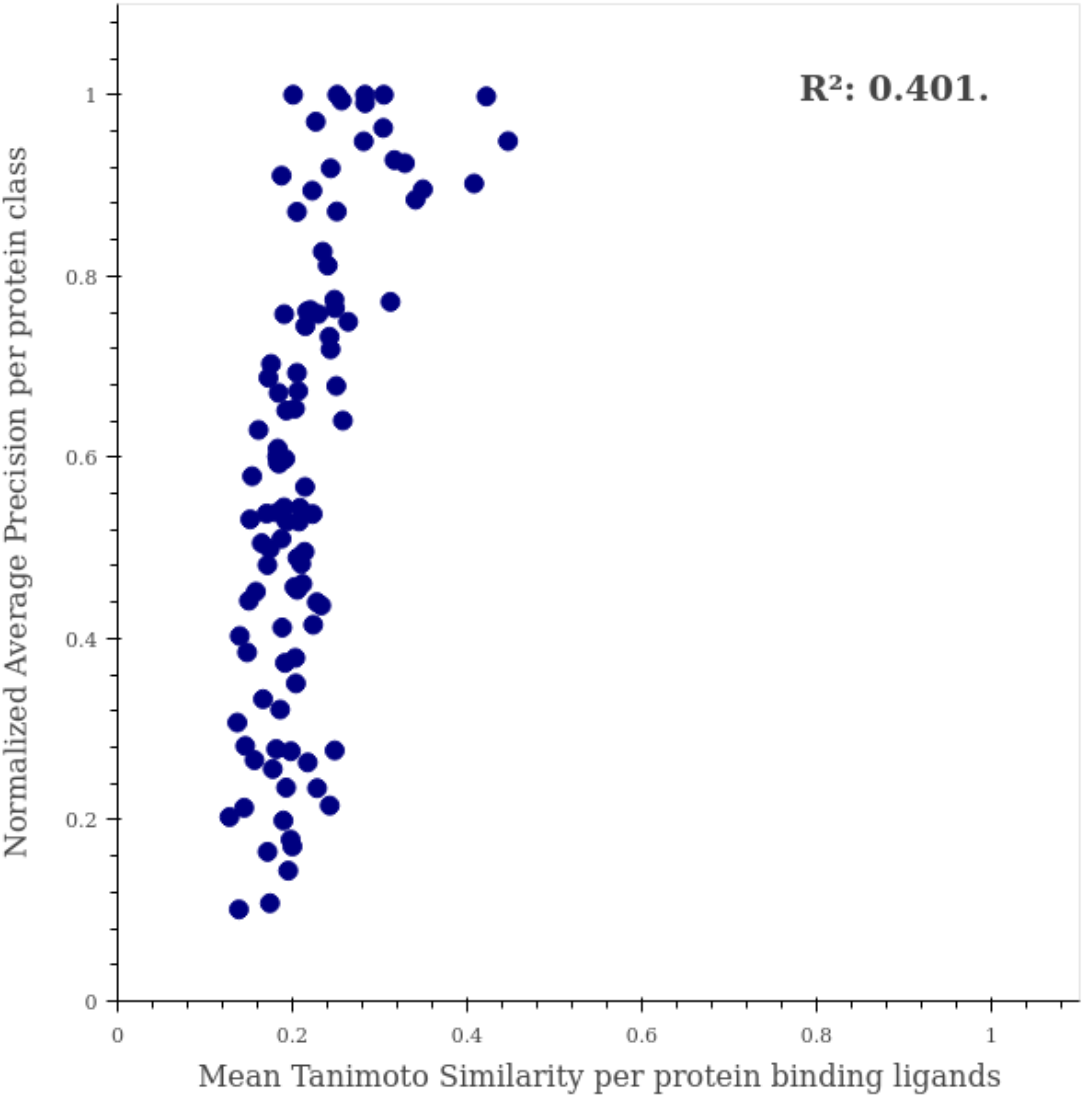
Correlation between Normalized AP and Tanimoto Similarity per protein. Performance per protein in PharmaNet is shown as a function of the averaged Tanimoto Coefficient in molecules of the same class. *R*^2^ is reported for estimating correlation between these two values.

Due to the increasingly growing interest of the pharmaceutical industry in antimicrobial treatments, and specifically in antivirals for enveloped viruses due to the possible impact on the ongoing coronavirus pandemic, we performed a screening for molecules active towards FPPS on a subset of the CHEMBL data set. The top-10 predicted molecules with a NAP of 100% are shown in Fig. 8. All molecules contain Nitrogen atoms, which have been reported previously to be essential in the interaction with the IPP active site of FPPS [68]. However, it can also be seen that none of the candidates has a phosphonic acid group, which we detected previously as a common moiety of active molecules towards FPPS. This unexpected outcome supports the idea that neural networks learn different fingerprints that those proposed and implemented by chemists for drug design for decades [11]. This shift on the molecule and fingerprint learning will enable the discovery of pharmaceuticals within a further representation space that consequently leads to significant structural differences. This, in turn, is likely to lead to revealing new and different mechanisms of action for known molecules. This is a very important characteristic for drug discovery since drugs with similar mechanisms of action are very likely to have the same drawbacks of current pharmaceuticals.

**Fig 8.**
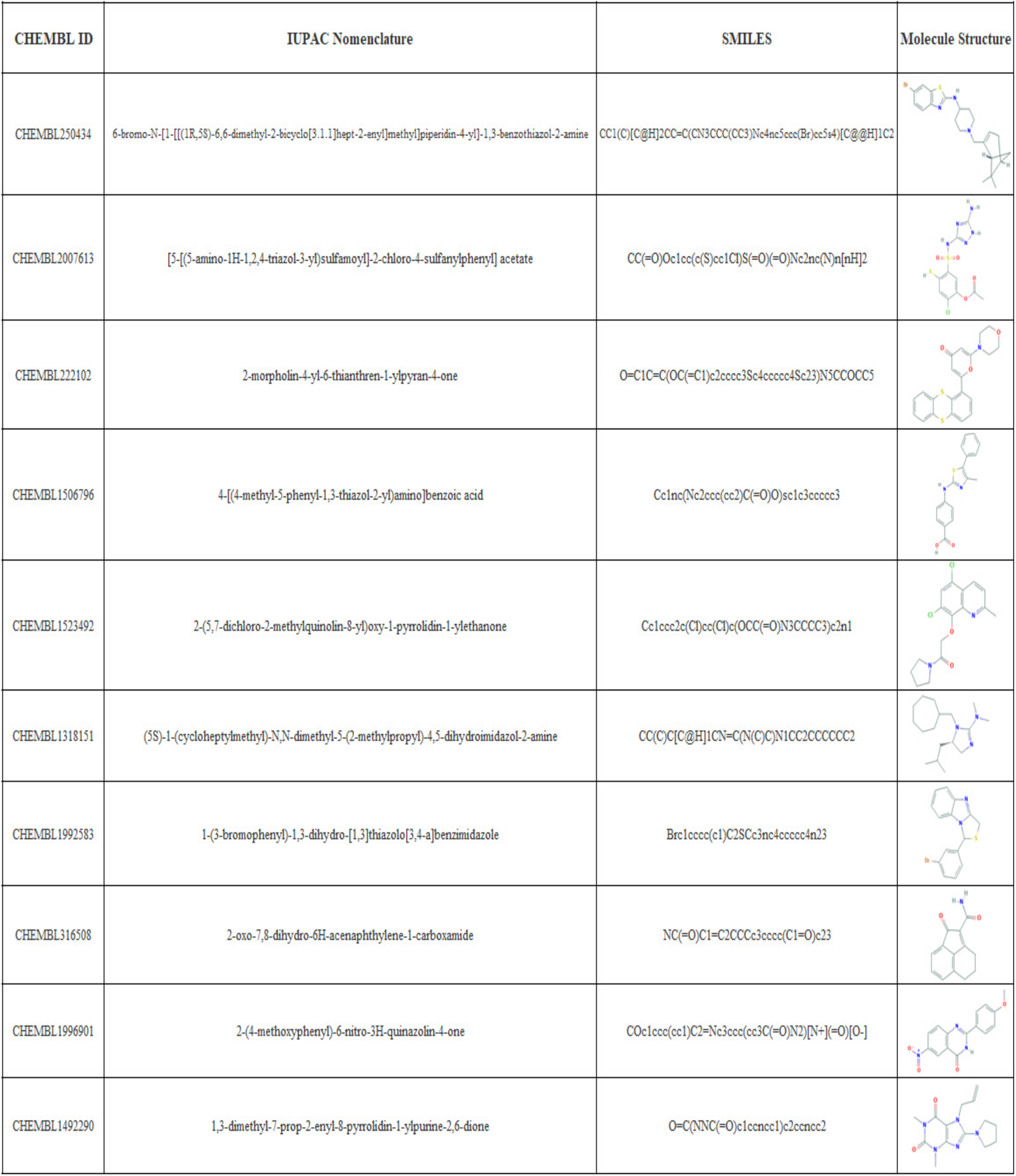
Top-10 predictions of PharmaNet towards FPPS on the CHEMBL subset data. CHEMBL ID, IUPAC name, SMILE and the molecule structure is given for the best 10 performing molecules in CHEMBL subset data towards FPPS target when predicting with PharmaNet.

To further corroborate the performance of our method, the top-10 candidates were analyzed via molecular docking and compared with Zolonadrate, a molecule reported to induce inhibition of FPPS (Table 3)[68]. All of our candidates showed binding energies lower than Zolonadrate’s and consequently, they should exhibit better affinity towards FPPS than Zolonadrate. This suggests a potential pharmacological use of the identified molecules and therefore the need for further toxicity studies. The interactions and binding energies with FPPS for the top-3 candidates and Zolonadrate are shown in Fig. 9.

**Table 3.** Binding Energies for top-10 candidates and inhibitor. Inhibitor in bold.

| CHEMBL ID | Binding Energy |
| --- | --- |
| CHEMBL250434 | -10.04 |
| CHEMBL2007613 | -9.56 |
| CHEMBL222102 | -8.48 |
| CHEMBL1506796 | -7.82 |
| CHEMBL1523492 | -7.60 |
| CHEMBL1318151 | -7.49 |
| CHEMBL1992583 | -7.43 |
| CHEMBL316508 | -6.56 |
| CHEMBL1996901 | -6.49 |
| CHEMBL1492290 | -5.94 |
| **Zolonadrate** | **-5.92** |

**Fig 9.**
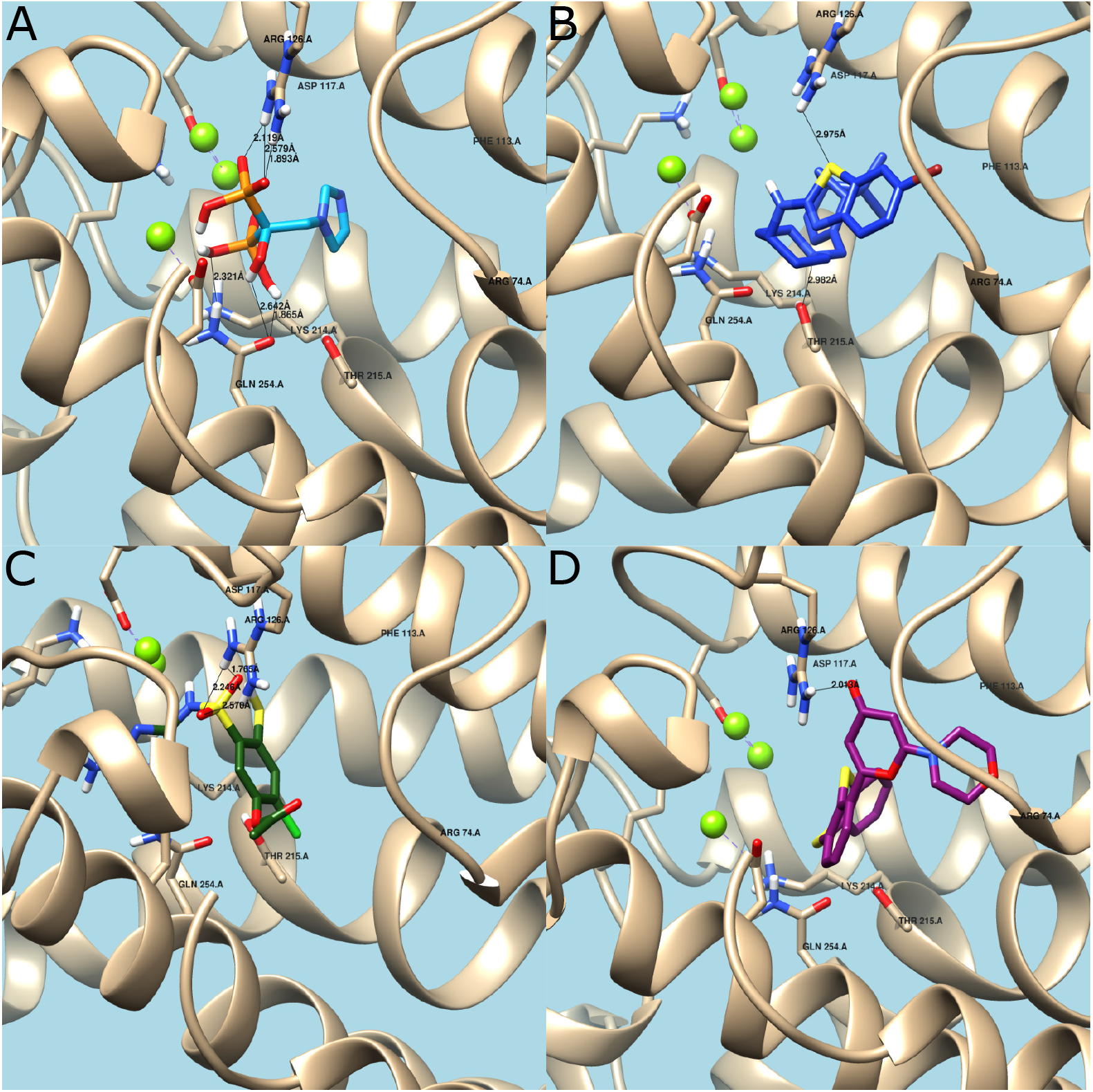
Molecular docking. (A) Zoledronate’s interaction with active site. Hydrogen bonds with Arg126 and Gln254. Binding Energy = −5.92. (B) CHEMBL250434’s interaction with active site. Hydrogen bonds with Arg126 and Thr215. Binding Energy = −10.04. (C) CHEMBL2007613’s interaction with active site. Hydrogen bonds with Arg126. Binding Energy = −9.56. (D) CHEMBL222102’s interaction with active site. Hydrogen bonds with Arg126. Binding Energy = −8.48.

The obtained binding energy for Zolonadrate agrees well with that reported elsewhere [55]. In this case (Fig. 8A), the prevalent interaction is by means of hydrogen bonding between the hydroxyl groups of Zolonadrate and the amine groups of the residues Gln254 and Arg126 [69]. Additional hydrogen bonding takes place between the protonated nitrogen atom of the heterocyclic ring in the side chain of Zolonadrate and the conserved main-chain carbonyl oxygen of Lys214 and the hydroxyl group of the Thr215. This stabilization mechanism resembles that of a carbocation intermediate [68].

CHEMBL250434 (6-bromo-N-[1-[[(1R,5S)-6,6-dimethyl-2-bicyclo[3.1.1]hept-2-enyl]methyl]piperidin-4-yl]-1,3-benzothiazol-2-amine) (Fig. 9B) has two heterocyclic rings in its structure, which could protonate and interact strongly with the protein at residues Lys214, Arg126, and Thr215 via hydrogen bonding. This is in contrast with Zoledronate, which has only one site for interaction. CHEMBL2007613 (5-[(5-Amino-4H-1,2,4-triazol-3-yl)amino]sulfonyl-2-chloro-4-mercaptophenyl acetate) (Fig. 9C) has one heterocyclic ring with Nitrogen atoms that can interact strongly with Lys214 and Arg216 and much less with Thr215. This is most likely the reason for the lower binding energy compared to CHEMBL250434 (6-bromo-N-[1-[[(1R,5S)-6,6-dimethyl-2-bicyclo[3.1.1]hept-2-enyl]methyl]piperidin-4-yl]-1,3-benzothiazol-2-amine). Finally, CHEMBL222102 (2-morpholin-4-yl-6-thianthren-1-ylpyran-4-one) (Fig. 9D) has one heterocyclic ring with only one Nitrogen atom capable of forming hydrogen bonds primarily with Arg126 residues, which explains the lowest biding energy of the analyzed set of molecules.

After corroborating the interaction through molecular docking, the top-3 candidates were analyzed with the aid of the online servers GUSAR, for cytotoxicity, and DIGEP-Pred for genotoxicity. Table 4 shows the toxicity label for the three compounds after administration via four different routes. Toxicity is categorized in a relative scale that goes from 1 to 5, with 1 for absence of toxicity and 5 the highest toxicity [57]. According to our findings, the only molecule with the potential for IV administration at the clinical level is CHEMBL2007613 (5-[(5-Amino-4H-1,2,4-triazol-3-yl)amino]sulfonyl −2-chloro-4-mercaptophenyl acetate).

**Table 4.** Citotoxicity predicted by GUSAR. NA: Non-Applicable to predictor domain.

| CHEMBL ID | IP (Intraperitoneal) | IV (Intravenous) | Oral | SC (Subcutaneous) |
| --- | --- | --- | --- | --- |
| CHEMBL250434 | Class 4 | Class 4 | Class 4 | NA |
| CHEMBL2007613 | NA | Non-Toxic | NA | NA |
| CHEMBL222102 | Class 4 | Class 4 | Class 4 | NA |

Regarding gene tocixity, CHEMBL2007613 upregulates the expression of the *PCDH17* gene, which encodes for a protein that contains six extracellular cadherin domains, a transmembrane domain, and a cytoplasmic tail, which makes it different from classical cadherins [70]. This gene has shown varying expression levels under some viral infections, which change depending on the type of virus [71]. Moreover, no reports are available discussing the downregulation of any other genes.

CHEMBL2007613 5-[(5-Amino-4H-1,2,4-triazol-3-yl)amino]sulfonyl-2-chloro-4-mercaptophenyl acetate (PubChem CID:380934) was established as non-toxic for IV administration routes and upregulates the *PCDH17* gene, which has been related to viral infections. Furthermore, this molecule is purchasable at the chemical companies ChemTik and ZINC. Given the above, we propose CHEMBL2007613 as a potential antiviral drug, for enveloped viruses, such as SARS-CoV-2.

## Conclusion

One important challenge in modern drug discovery is to accelerate the search for new and more potent therapeutic molecules but in a more assertive manner as thus far, conventional approaches are extremely costly, time consuming and largely ineffective. Pharmaceutical companies have spent billions of dollars in development but only a small percentage of candidates have made it to the market. Novel artificial intelligence algorithms provide an alternative route for a more comprehensive search of candidates in already available and large databases of pharmaceutical compounds. Also, those algorithms reduce the number of experiments needed *in vitro* and *in vivo*, given that only the most promising candidates are further analyze. We implemented this approach here and put forward PharmaNet, a deep learning architecture for predicting binding of a molecule to possible protein target receptors. PharmaNet’s algorithm represents the 2D structural information of a molecule as a molecular image and process it with modern computer vision recognition techniques. Our architecture is trained end-to-end and consists of a convolutional encoder processing phase followed by an RNN. This approach allows multiclass classification with a single model. The conventional metric for this type of task has been the Receiver Operating Characteristic curve (ROC-AUC); however, we propose that a more accurate metric is the area under the Normalized Average Precision (NAP) curve. Under this framework, PharmaNet outperforms the state-of-the-art algorithm by one order of magnitude (from 1% to 65.5%) in the AD dataset, and has a perfect performance in identifying the active molecules for 3 receptors of the 102 targets: CXCR4, FPPS and KITH. We selected FPPS as target to apply our model in the search of active molecules within the large database of pharmacological molecules, ChEMBL. We chose the 10 best candidate molecules to investigate into detail interactions with FPSS via molecular docking. We also conducted an *in silico* evaluation of toxicity with the aid of online servers. We found that the compound identified with the ID CHEMBL2007613, i.e., (5-[(5-Amino-4H-1,2,4-triazol-3-yl)amino]sulfonyl-2-chloro-4-mercaptophenyl acetate) exhibits potential antiviral activity, which needs to be corroborated *in vitro*. We expect that our algorithm opens new opportunities for the rediscovery and repurpose of pharmacological compounds that otherwise might be disregarded in importance by the pharmaceutical industry. Moreover, we are currently exploring its potential in the reverse problem, i.e., searching for multiple receptor targets for molecules with certain physicochemical properties.

## Acknowledgments

We sincerely thank authors from [25] for kindly giving us their predictions in order to make a more direct comparison of both methods.

